# Antibody Cocktail Exhibits Broad Neutralization against SARS-CoV-2 and SARS-CoV-2 variants

**DOI:** 10.1101/2021.04.16.440083

**Authors:** Yuanyuan Qu, Xueyan Zhang, Meiyu Wang, Lina Sun, Yongzhong Jiang, Cheng Li, Wei Wu, Zhen Chen, Qiangling Yin, Xiaolin Jiang, Yang Liu, Chuan Li, Jiandong Li, Tianlei Ying, Dexin Li, Faxian Zhan, Youchun Wang, Wuxiang Guan, Shiwen Wang, Mifang Liang

## Abstract

Severe acute respiratory syndrome coronavirus 2 (SARS-CoV-2) has precipitated multiple variants resistant to therapeutic antibodies. In this study, 12 high-affinity antibodies were generated from convalescent donors in early outbreaks using immune antibody phage display libraries. Of them, two RBD-binding antibodies (F61 and H121) showed high affinity neutralization against SARS-CoV-2, whereas three S2-target antibodies failed to neutralize SARS-CoV-2. Following structure analysis, F61 identified a linear epitope located in residues G446 -S494, which overlapped with angiotensin-converting enzyme 2 (ACE2) binding sites, while H121 recognized a conformational epitope located on the side face of RBD, outside from ACE2 binding domain. Hence the cocktail of the two antibodies achieved better performance of neutralization to SARS-CoV-2. Importantly, F61 and H121 exhibited efficient neutralizing activity against variants B.1.1.7 and B.1.351, those showed immune escape. Efficient neutralization of F61 and H121 against multiple mutations within RBD revealed a broad neutralizing activity against SARS-CoV-2 variants, which mitigated the risk of viral escape. Our findings defined the basis of therapeutic cocktails of F61 and H121 with broad neutralization and delivered a guideline for the current and future vaccine design, therapeutic antibody development, and antigen diagnosis of SARS-CoV-2 and its novel variants.

## Introduction

Severe acute respiratory syndrome coronavirus 2 (SARS-CoV-2) is acknowledged as the novel coronavirus that causes the global pandemic of COVID-19 (Andersen *et al*., 2020). Up to 28 February 2021, over 100 million confirmed cases have been reported worldwide (World Health Organization). SARS-CoV-2 grouped to the betacoronavirus genus (Wu *et al*., 2020) is proved to share about 80% sequence identity to SARS-CoV and target the same cellular receptor, angiotensin-converting enzyme 2 (ACE2) (Daniel *et al*., 2020). ACE2 directly binds to SARS-CoV-2 spike (S) protein which is consisted of S1 subunit and S2 subunit (Daniel *et al*., 2020).

To date, a variety of neutralizing antibodies against SARS-CoV-2 S protein have been generated. Potent neutralizing often found to target S1 subunit (Hwang *et al*., 2006), which consists of the N-terminal domain (NTD) and the receptor-binding domain (RBD). However, neutralizing antibodies targeting the S2 subunit still need to be discovered (Liu *et al*., 2020; Wec *et al*., 2020). The NTD-specific monoclonal antibodies (mAbs) target a patch remote from RBD (Chi *et al*., 2020; Liu *et al*., 2020). RBD-specific mAbs are divided into four main classes (Barnes *et al*., 2020b). Antibodies grouped in class one and class two, such as CB6 (Shi *et al*., 2020) and P2B-2F6 (Ge *et al*., 2021), are found with high potencies and overlapped with the receptor-binding motif (RBM) on RBD (Ju et al., 2020; Shi et al., 2020; Wu et al., 2020). These mAbs are dominant in convalescent serum (Piccoli *et al*., 2020). Antibodies in the third and fourth class, like S309 (Pinto *et al*., 2020) and CR3022 (Piccoli et *al*., 2020; Xiang *et al*., 2020), are positioned detached from the RBM (Starr *et al*., 2020)..

The rapid global spread and transmission of SARS-CoV-2 are hypothesized to provide the virus with substantial opportunities for the natural selection of favorable mutations, many of which involved modification of S protein. The D614G mutation in the S protein enhances viral transmission and overtakes the prime strain of SARS-CoV-2 (Zhang *et al*., 2020; Li et *al*., 2021). The recent emerging variants of concern observed in the United Kingdom(B.1.1.7 with mutations N501Y, A570D and del69/70), South Africa (B.1.351 with mutations K417N, E484K and N501Y), and Brazil (P.1 and P.2 with mutations K417T, E484K and N501Y) (Long *et al*., 2021) initially respond more tightly to ACE2 and appear to be more infectious to human (Laffeber *et al*., 2021; Tian *et al*., 2021). More severely, B.1.351 and P.1 are resistant to convalescent plasma, vaccine sera and multiple neutralizing mAbs (Hoffmann *et al*., 2021; Widera *et al*., 2021). Variants B.1.141 and B.1.258 with mutation N439K increase spike affinity for ACE2 and confer resistance to several mAbs (Thomson *et al*., 2021). American variants(B.1.429 and B.1.427)containing L452R(Long *et al*., 2021) show refractory to mAbs as well. SARS-CoV-2 variants isolated from minks and mouse harboring mutations G261D, A262S, L452M, Y453F, F486L, Q498H and N501T may cause potential cross-species transmission that worth closely monitor (Thomson *et al*., 2021; Yao *et al*., 2021). Thus, it is essential to develop antibodies with broad-spectrum activities against SARS-CoV-2 and SARS-CoV-2 variants.

A variety of neutralizing antibodies against SARS-CoV-2 have entered clinical trials. However, the virus may persist due to mutations, especially mutations on the S protein, leading to dropping neutralizing activity and hence efficacy from these neutralizing antibodies in the longer term (Li *et al*., 2020). Therefore, it is essential to develop various neutralizing antibodies against different epitopes. Besides, novel delivery strategies of antibodies, such as antibody inhalation treatment, would be encouraged and benefit from the convenience and widely applied during COVID-19 prevention.

Here we reported 12 mAbs screened with purified SARS-CoV-2 RBD, S1 and S2 from three COVID-19 convalescent patients by phage antibody library technique. Then we characterized their affinity, neutralizing activity and binding sites. We also evaluated the neutralizing activity of screened mAbs to SARS-CoV-2 variant. Additionally we selected two RBD-specific antibodies (F61 and H121) with high neutralizing activity and high affinity to investigate interaction between antibodies and RBD via computer simulation. Our research provided a theoretical basis for the development of therapeutic antibodies.

## Materials and Methods

### Cells and Viruses

Cell lines (HEK293T and VeroE6 cells) were initially acquired from the American Type Culture Collection (ATCC; USA). EXPi293F cells were purchased from Life Technologies, USA. They were cultured at 37 °C under 5% CO_2_ in Dulbecco’s modified Eagle’s medium (DMEM; Life Technologies, USA) supplemented with 10% heat-inactivated fetal bovine serum (FBS; Life Technologies, USA) and 1% penicillin/streptomycin (Life Technologies, USA) or in EXPI293 expression medium (Life Technologies, USA). Cells were passaged every two days and digested with 0.05% trypsin-EDTA. Pseudovirus of SARS-CoV-2 (GenBank: MN908947) and SARS-CoV-2 variants were obtained from National Institutes for Food and Drug Control. The authentic SARS-CoV-2 (GenBank: MN908947) and SARS-CoV-2 variants were obtained from Wuhan Institute of Virology. All work with infectious SARS-CoV-2 was performed in Institutional Biosafety Committee approved BSL3 facilities using appropriate positive pressure air respirators and protective equipment.

### Construction and Screening of Human Antibody Phage Display Library

The phage display library procedures in the vector pComb 3H followed the methods described previously (Kashyap *et al*., 2008). Briefly, lymphocytes were isolated from three convalescent donors in early outbreaks which were selected by Enzyme-Linked Immunosorbent (ELISA) assays and colloidal gold test (INNOVITA, CHN). Total cellular mRNA was extracted using the RNeasy Mini kit (Qiagen, GER), and cDNA was synthesized with primer oligo (dT) using Transcriptor High Fidelity cDNA Synthesis kit (Roche, SUI). PCR amplification was then performed using FastStrat High Fidelity PCR System (Roche, SUI). The light and heavy chain genes were amplified from the cDNA by PCR using the primer pairs from VK, VL and VH gene families, then cloned into the vector pComb 3H(Barbas *et al*., 1991). The library’s initial diversity was evaluated and assured by sequencing of randomly picked clones for each step of library construction and the complexity of the library was then calculated. The final yielded antibody libraries were panned and screened with purified SARS-CoV-2 RBD protein, S1 protein and S2 protein (Jiangsu East-Mab Biomedical Technology, CHN) following the standard panning procedure(Barbas and Burton, 1996).

### Production of monoclonal antibody

For recombinant human mAb production, the cDNA encoding mAb variable regions of the heavy and light chains were cloned into expression plasmids containing the human IgG1 heavy chain and Ig kappa or lambda light chain constant regions, respectively. Recombinant mAbs were then produced in EXPi293F cells (Life Technologies, USA) by transfecting pairs of the IgG1 heavy and light chain expression plasmids. Human antibodies purified by Protein-G (GE Healthcare, USA) affinity chromatography were stored at -80°C until use.

### Enzyme-Linked Immunosorbent (ELISA) Assays and non-competitive ELISA assay

ELISA plates were coated with SARS-CoV-2 RBD protein, S1 protein, S2 protein, S protein trimer and mutant S1 protein (Jiang-su East-Mab Biomedical Technology, CHN) at 4 °C overnight. Following washing with PBST, serial dilutions of testing antibodies start at 1μg/ml or serial dilutions of plasma start at 1:100 were added to each well and incubated at 37 ° C for 30min. After washing with PBST, horseradish peroxidase (HRP)-conjugated anti-human IgG antibody (Sigma, USA) was added at the dilution of 1:20000 and incubated at 37°C for 30min. The absorbance was detected at 450nm. The data was analyzed using GraphPad Prism 8.0.

### Surface plasmon resonance (SPR) assay

Purified antibodies targeting S1 were quantified with SPR assay using the BIAcore 8000 system (GE Healthcare, USA) carried out at 25°C in single-cycle mode. Purified SARS-CoV-2 S1 diluted in 10 mM sodium acetate buffer (PH 5.5) was immobilized to CM5 sensor chip by amine coupling reaction. Serially diluted antibodies were injected with a rate of 30 ml/min in sequence. The equilibrium dissociation constants (binding affinity, Kd) for each antibody were calculated using Biacore 8000 Evaluation Software.

### Virus neutralization assay

The virus neutralization assay with pseudoviruses was conducted as described previously (Nie *et al*., 2020). Briefly, serially diluted antibodies were added into 96-well plates. After that, 50 µl pseudoviruses were added to the plates, followed by incubation at 37°C for one hour. Afterward, HuH-7 cells were added into the plates (2×10^4^ cells/100 µl per well), followed by 24h incubation at 37°C in a humidified atmosphere with 5% CO_2_. Chemiluminescence detection was performed straight after, and the Reed-Muench method was used to calculate the virus neutralization titer. The half-maximal inhibitory concentrations (IC50) were determined using 4-parameter logistic regression (GraphPad Prism version 8).

Authentic SARS-CoV-2 was used in the plaque reduction neutralization test (PRNT). In brief, the mAbs were trifold serially diluted in culture medium and mixed with SARS-CoV-2 (200 PFU) for one hour. Mixtures were then transferred to 24-well plates seeded with Vero E6 cells and allowed absorption for 1 h at 37 °C. Inoculums were then removed before adding the overlay media (100 μl MEM containing 1 % carboxymethylcellulose, CMC). The plates were then incubated at 37 °C for 96 h. Cells were fixed with 4% paraformaldehyde solution for one day, and overlays were removed. Cells were incubated with 1% crystal violet for five minutes at room temperature. The half-maximal inhibitory concentrations (IC50) were determined using 4-parameter logistic regression (GraphPad Prism version 8).

### Fluorescence-activated cell sorting (FACS) assay

SARS-CoV-2 S protein-expressing plasmids were transfected into HEK293T cells using Lipofectamine 3000 (Invitrogen, USA). 24 h after transfection, cells were suspended and washed with PBS twice. Then the cells were incubated with 20 μg/ml mAbs or isotype IgG mAb of hepatitis b virus (HBV), at room temperature for 1 h, followed by further incubation with anti-human IgG FITC-conjugated antibody (Sigma, USA). The cells were analyzed using FACSAria II(BD, USA). All of these data were analyzed using FlowJo.

The block assay was assessed by FACS. HEK293T cells were transiently transfected with the ACE2 expression plasmid for 24 h. The mouse-Fc tag Fusion protein of SARS-CoV-2 RBD (RBD-mFC) (Jiang-su East-Mab Biomedical Technology, CHN) at a concentration of 2 µg/ml was mixed with the mAbs or isotype IgG at a molar ratio of 1:10 and incubated at 4 °C for 1 h. Then mixtures were added to 2.5 × 10^5^ HEK293T cells expressing ACE2 and incubated at 4 °C for another hour. Then cells were stained with anti-mouse IgG Taxes red-conjugated antibody and anti-human IgG FITC-conjugated antibody (Sigma, USA) for another 30 min then analyzed by FACSAria II (BD, USA).

### Competition ELISA assay

Plates were coated with SARS-CoV-2 RBD (Jiang-su East-Mab Biomedical Technology, CHN) at 4 °C overnight. Two-fold serial dilutions antibodies were added to the wells, and plates were incubated for one hour at 37°C. Then plates were incubated with HRP-conjugated mAb (diluted 1:2000) (Wantai BioPharm, CHN) for 30min at 37°C. HRP activity was measured at 450 nm. Paired antibodies with a value less than 20% were defined as noncompeting. Antibodies were deemed to compete for the same epitopes if the value was calculated greater than 60%. Otherwise, if the value was found between 20% and 60%, the antibody pairs were considered partially overlapping. The percent of binding inhibition of labeled antibodies was calculated according to the formula below:

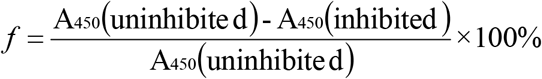

### Molecular Modeling and Docking of the antibodies to RBD

The computational simulation was carried out by Discovery studio 2.0 (Accelrys, San Diego, CA)(Kaushik and Sowdhamini, 2011). A suitable template was obtained through a BLAST search of the Protein Databank (PDB). The homology modeling of mAbs was performed using DS Homology Modeling protocol, and the 3D model of antibody was optimized using Antibody loop refinement protocol. The models were validated by Ramachandran plots. Protein-protein docking of RBD and mAbs was performed using the ZDOCK and RDOCK programs by specifying the variable region’s antibody residues on the binding interface. RDOCK refinement was performed on the top 100 poses of the filtered ZDOCK output and applied scoring function to each docked structure for best binding models.

## Result

### Generation and screening of Antibodies Against SARS-CoV-2

To isolate mAbs, we collected plasma and peripheral blood mononuclear cells (PBMCs) from 15 confirmed COVID-19 convalescent patients in Hubei and Shandong Provence. We evaluated antibodies titer in plasma to SARS-CoV-2 N protein and different fragments of S protein, including RBD, S1, and S2 with ELISA (Fig. 1A) and colloidal gold test (data not shown). The plasma from donors 2, 10 and 11 showed higher IgG titer against RBD, S1 and S2. Thus they were chosen for library construction by the pComb 3H vector system. The library was established with a complexity of 1×10^8^ estimated independent clones and 100% Fab genes diversity after sequencing confirmation. Single clone screening was performed with ELISA, and a total of 274 positive monoclonal were identified (Fig. 1B). Based on sequencing and ELISA results, four unique clones from RBD, five from S1, and three from S2 were chosen as the candidates for further interrogation.

**Figure 1.**
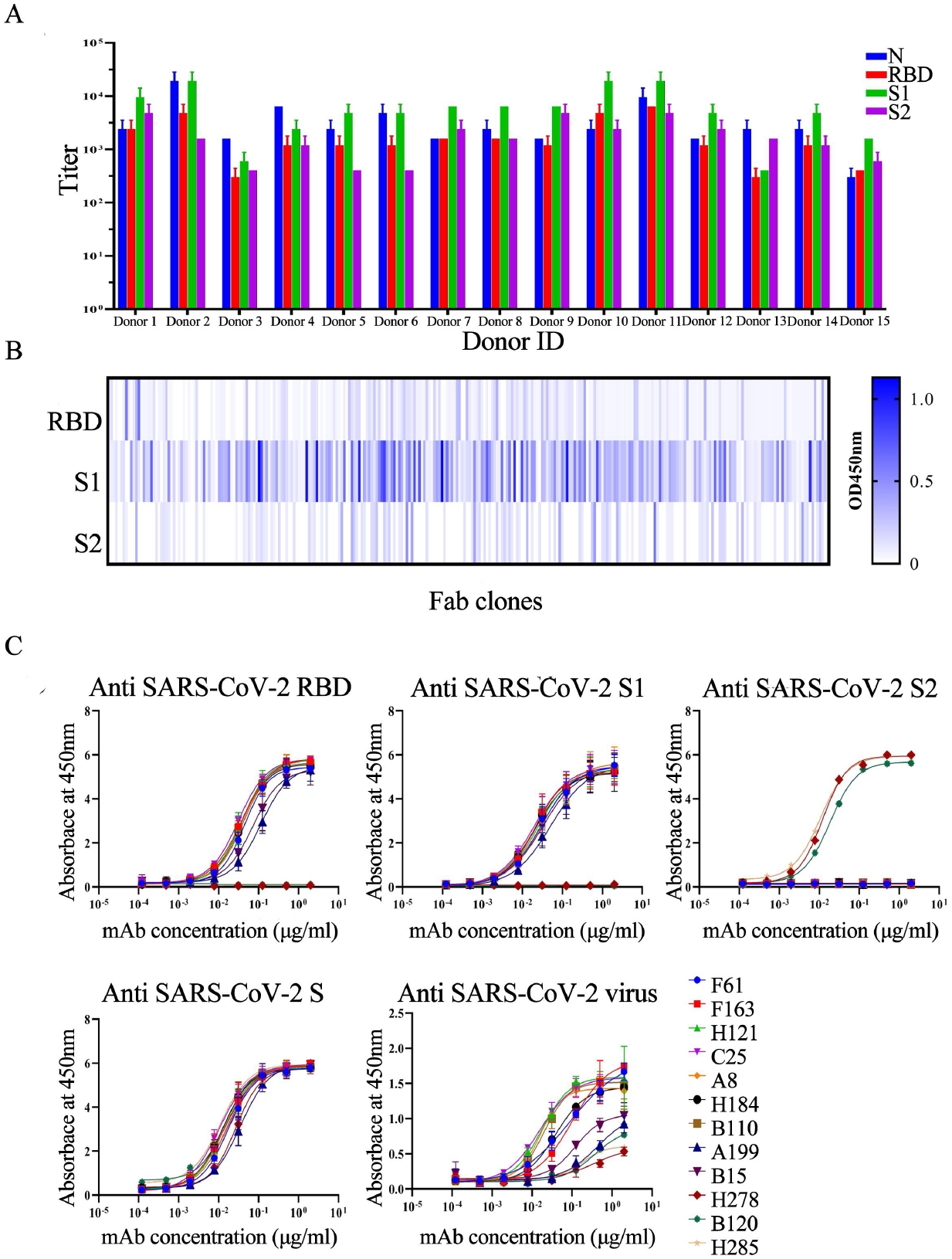
Generation and screening of antibodies from SARS-CoV-2 convalescent patients. (A) Antibodies titer in the plasma of SARS-CoV-2 convalescent patients to SARS-CoV-2 N protein and different fragments of SARS-CoV-2 S protein (RBD, S1, and S2). Experiments were performed in duplicate and the error bars denote ± SD, n = 2. (B) Heat-maps of Fab clones against RBD (n = 288), S1 protein (n = 288) and S2 protein (n = 288). Each lattice represented a Fab clone. (C) Binding specificity of the 12 candidate IgGs. The binding to different spike proteins (RBD, S1, S2, S protein trimer, and virion) was determined by ELISA Experiments were performed in duplicate, and the error bars denote ± SD, n = 2.

In order to analysis the characteristics of selected Fab antibodies, we recloned the Fab antibodies into the IgG1 format. We further determined the binding specificity of the 12 candidate IgGs with purified SARS-CoV-2 virion and different S protein fragments (S1, S2, RBD and S protein trimer) utilizing ELISA. All 12 antibodies were able to recognize purified SARS-CoV-2 virion and S protein trimer. However, the binding strength of 12 antibodies to purified SARS-CoV-2 virion varied. A199, B15, H278, B120 and H285 had a weak affinity to virion. Nine antibodies (F61, F163, B15, H121, C25, A8, H184, B110 and A199) screened with purified RBD and S1 were all positive to RBD. The rest three of them (H278, B120 and H285) were found attached to S2 but not S1 nor RBD (Fig. 1C). However, NTD specific mAbs were not screened and identified.

### Characterizing the binding profile of 12 candidate antibodies

The specificity of candidate SARS-CoV-2 specific IgGs were evaluated utilizing FACS. All candidate IgGs (labeled by FITC) showed positive on the surface of HEK293 T cells expressing SARS-CoV-2 S protein. In contrast, HBV mAb, as a negative control, demonstrated no interaction with S protein (Fig. 2A and Fig.S1 A). Thus, all tested mAbs were suggested binding specifically to SARS-CoV-2 S protein.

**Figure 2.**
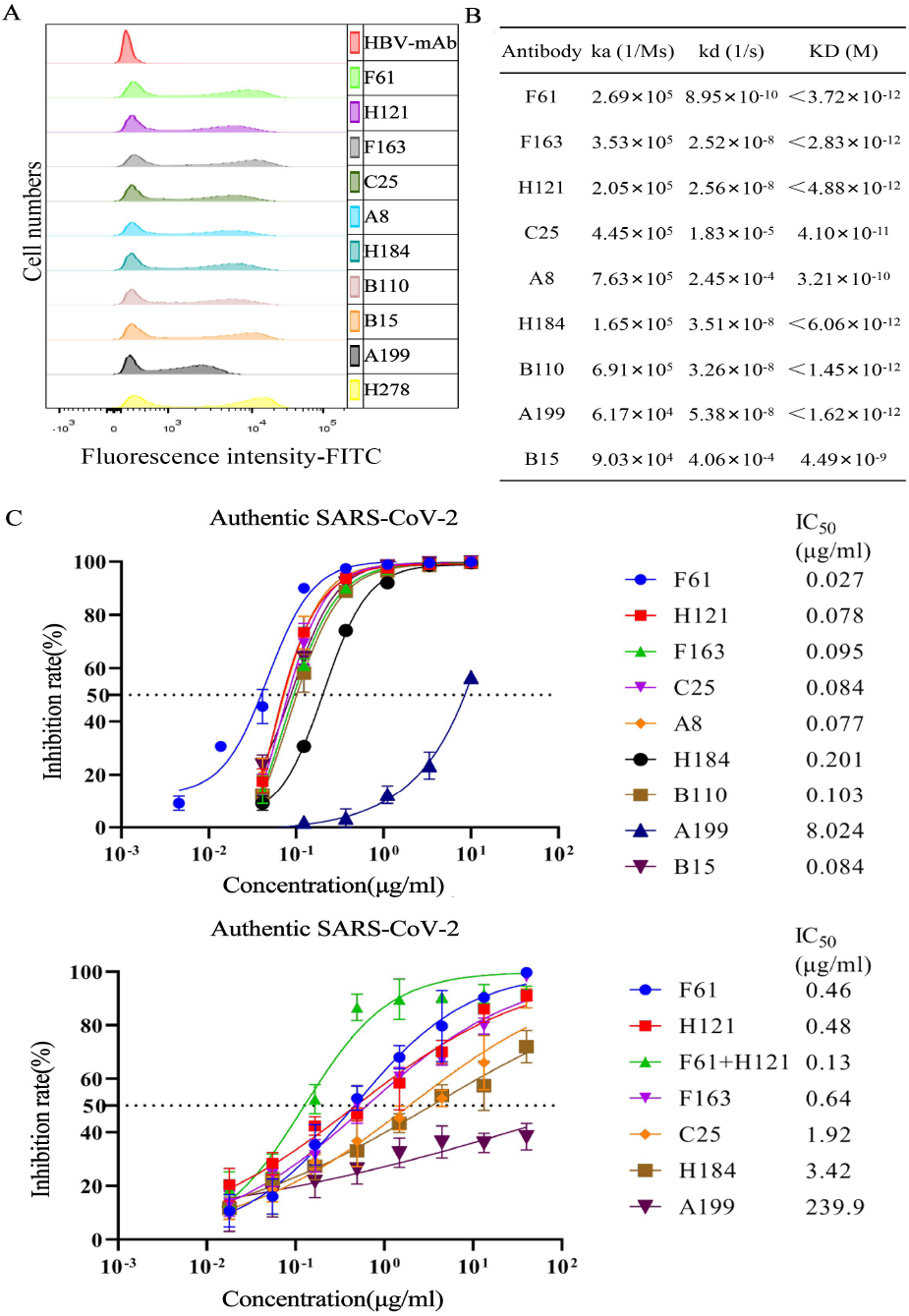
Characterizing the binding profile of candidate IgGs. (A) The specificity of SARS-CoV-2 specific IgGs detected by FACS. HEK 293T cells expressing SARS-CoV-2 S protein were incubated with candidate mAbs or isotype IgG (HBV mAb) and then stained with anti-human IgG FITC-conjugated antibody. Fluorescence intensity(FITC) negative cells was less than 10^3^, and that of positive cells was around 10^4^. (B) The affinity of candidate IgGs.. The affinity between antibodies (F61, F163, B15, H121, C25, A8, H184, B110 and A199) and S1 were measured by BIAcore 8000 system. Non-competitive ELISA measured the affinity between mAbs(H278, B120 and H285) and S2. (C) Neutralizing activity of candidate IgGs against WT-SARS-CoV-2 pseudovirus and WT-authentic SARS-CoV-2. Experiments were performed in duplicate, and the error bars denote ± SD, n = 2. The dashed line indicated a 50% reduction in viral infection.

SPR assay were performed to evaluate the affinity of nine RBD-specific IgGs to S1 protein. F61, F163, H121, C25, H184, B110 and A199 showed a high affinity to S1 protein. The KD values ranged from 1.45×10^−12^M to 4.88×10^−12^M. In comparison, lower KD values were detected regarding B15 and A8 (Fig. 2B and Fig.S1 B). Non-competitive ELISA assay were performed to evaluate the affinity of H278, B120 and H285 against S2 protein. The KD values of H278, B120 and H285 were 2.15×10^−11^M, 2.6×10^−11^M, 2.74×10^−11^M, respectively(data not show)

Neutralizing capacity of 12 candidate IgGs were evaluated by authentic SARS-CoV-2 neutralization assay and pseudoviruses neutralization assay. F61, H121 and F163 exhibited high neutralizing capacity with IC50 of 0.46 ug/ml, 0.48 ug/ml, and 0.64 ug/ml to authentic SARS-CoV-2, and 0.027 ug/ml, 0.078 ug/ml, and 0.095 ug/ml to pseudoviruses, respectively. However, A199 exhibited low neutralizing capacity, which which suggested not all RBD-specific antibodies were neutralizing antibodies. S2-specific mAbs failed to neutralize SARS-CoV-2 (data not shown). Moreover, the cocktail of F61 and H121 exhibited a synergistic neutralization to authentic SARS-CoV-2 with the neutralizing capacity (0.13 ug/ml) increased four times compared to each one alone (Fig. 2C).

To further characterize nine RBD-specific IgGs, we then analyzed antigenic epitopes of nine RBD-specific antibodies by competitive ELISA. Antibodies were roughly classified into three groups by competition percentages. Group one contained F61, F163 and B15. Group two included H121, C25, A8, H184 and B110. Group one and group two were not competitive, which suggested they bound to different antigenic epitopes. A199 from group three had no competition with mAbs in group one yet a partial competition with those from group two. A199 bound to a non-neutralization epitope on RBD (Fig. 3A).

**Figure 3.**
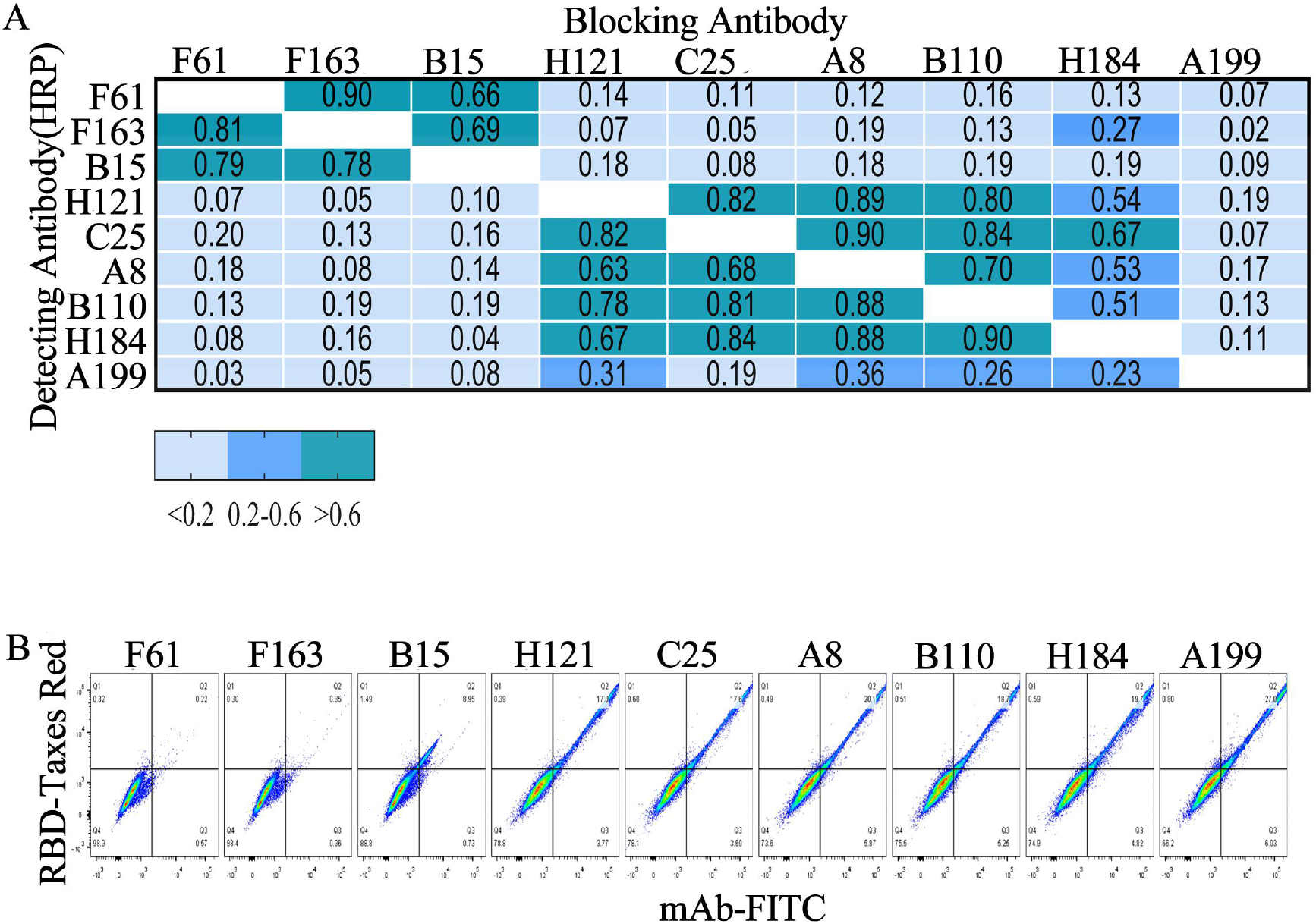
Analysis of antibodies epitopes by Competition ELISA assays and FACS. (A) Antigenic epitopes of nine RBD-specific IgGs were analyzed by competitive ELISA. Each lattice shows a competitive percentage. Values less than 0.20 indicated that the antibody has non-competitive epitopes, the value between 0.20 and 0.60 indicated intermediate binding sites, and values greater than 0.60 indicated that the antibody shares overlapping or tight epitopes. (B) ACE2 binding block assay by FACS. The mouse-Fc tag Fusion protein of SARS-CoV-2 RBD (RBD-mFC) was pre-incubated with nine RBD-specific IgGs or isotype IgG (HBV mAb) and then stained with HEK 293T cells expressing ACE2. Anti-human (Fc) FITC-conjugated antibody and Anti-mouse (Fc) Texas red-conjugated antibody were used as the secondary antibody. The X-axis represented the fluorescence intensity of human antibodies labeled by FITC, and the Y-axis represented the fluorescence intensity of RBD-mFC labeled by Taxes red.

The inhibitory effect of these antibodies on the RBD-ACE2 interaction was investigated by FACS using ACE2 expressing HEK293T cells. F61 and F163 from group one prevented RBD from binding to ACE2 with no fluorescence signal from the antibody (FITC) or RBD (Taxes red), which suggested they bound to an ACE2-competitive epitopes. B15 only partly blocked the binding of RBD to ACE2 due to its low affinity and neutralization capacity. In group two, H121, C25, A8, H184 and B110 failed to block the binding between RBD and ACE2 with double-positive cells to antibody and RBD. Their epitopes were far from ACE2-binding domain on RBD. As the neutralization capacity of antibodies decreased, more cells showed double-positive to antibody and RBD. The proportion of double-positive cells to H121, which suggested the highest neutralization capacity and affinity, was 17%. The proportion of double-positive cells to H184, one with lower neutralization capacity, was 19.7%. A199 was also double-positive to antibody and RBD. The neutralizing capacity of A199 was negligible, consistent with its poor performance in preventing the binding of ACE2 to RBD with the highest proportion of double-positive cells (Fig. 3B).

So far, we have obtained three types of antibodies, one of which is mainly represented by F61 and F163 to recognize ACE2 receptor epitopes with high neutralization activity. One is represented by H121 to identify epitopes which not overlap with ACE2 binding sites, but it has high neutralization activity. And the last one is A199 which bound to a non-overlapping epitope with ACE2 binding sites had no neutralization activity.

### Interaction between mAbs and RBD via Computer Modeling and Docking

F61 and H121 exhibited the high neutralization capacity and bound to different epitopes. They were excellent candidate of antibody-based drugs to SARS-CoV-2. To precisely delineate the interaction between antibody and antigen. We further investigated interaction between F61/ H121 and RBD via computer simulation. Crystal structures that shared over 90% sequence similarity with F61 and H121 were used as the antibody template to build the 3D-structure of the two antibodies. Meanwhile, crystal structure 7DK3 (PDB) (Daniel et al., 2020) of SARS-CoV-2 RBD was used in the ZDOCK program and RDOCK program.

The outcomes demonstrated that F61 and H121 bind to diversified regions on RBD. F61 identified a linear epitope ranging from G446 to S494 within the RBM region involving 23 residues on RBD (Fig. 4A). The predicted binding sites indicated both the light and the heavy chain of F61. In specific, hydrogen-bond (H-bond) could be formed upon approaching RBD’s P479, C480,N481 and F486 with D108, N37 and S109 on the F61’s light chain, as well as F490 and L492 with G109 and R36, E484 with Y38 and Y114 on the heavy chain (Fig.4B, upper panel). In comparison, H121 recognized a conformational epitope located remote from the RBM region and mainly contributed by the heavy chain (Fig. 4A). Specifically, R355, G381 and L517 on RBD formed H-bond with S11A, N59 and Y109, L518 and A520 to Y37 on the heavy chain. R357 bond to Y38 on the light chain (Fig.4B, lower panel).

**Figure 4.**
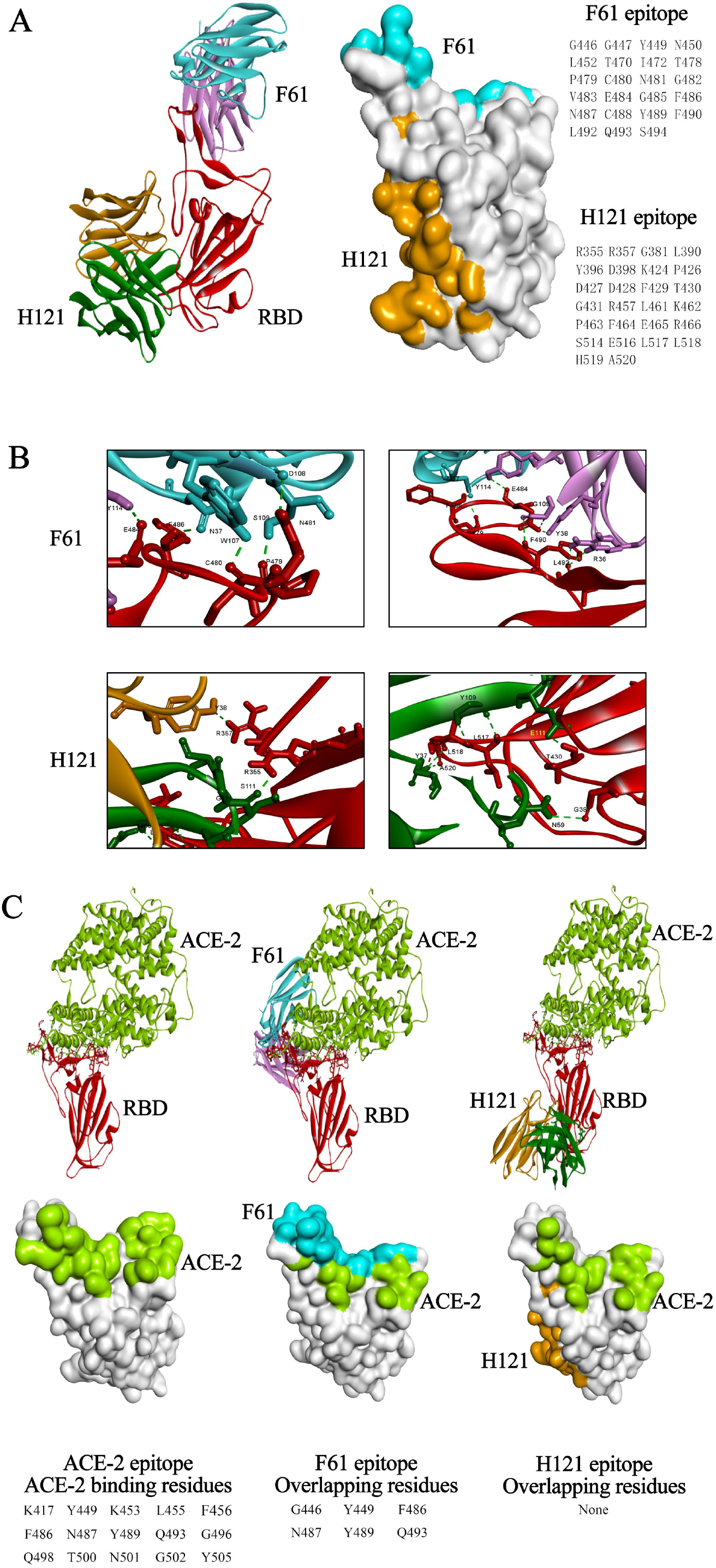
Computer docking (ZDOCK) structure between F61/H121 and SARS-CoV-2 RBD. (A)ZDOCK structure of the RBD and antibodies complex was shown on the left. RBD was in red. F61 colored blue and pink. H121 was in orange and green. The complex of two antibodies and RBD were superimposed to demonstrate their relative positions and orientations. The footprint of F61 and H121 on RBD was shown in the middle. Blue and orange represent the footprint of F61 and H121, respectively. Binding residues were listed on the right. (B) The predicted Hydrogen-bond (H-bond) of F61 and H121. Green dashed lines indicated H-bond. The H-bond of F61 was shown on the upper panel. The H-bond of H121 was shown on the lower panel. (C) Epitopes were overlapping between the two antibodies and ACE2. The interaction between ACE2 and RBD was shown on the left. RBD was in red. ACE2 colored bright green. The interactions between ACE2, RBD and two antibodies were shown on the middle (F61) and left (H121). Color settings were consistent with those mentioned above. Overlapping residues between each of the antibodies and ACE2 were listed at the bottom.

The antibody-RBD complex structure was subsequently aligned with the ACE2-RBD complex based on the RBD sequence (Fig.4C upper panel). Six residues within the F61 epitopes overlapped the ACE2-binding sites on the RBD. H121 positioned further away from the ACE2-binding sites and hence showed no competitiveness in previous assays (Fig.4C lower panel).

### Determination of the effects of natural mutations in S protein on the sensitivity of candidate antibodies

All S protein mutations (reported in the GISAID database up to 19 January 2021) were retrieved for analysis, with 333251 sequences selected. Amino acid replacements, insertions, and deletions with a frequency exceeding 0.1% were focused on our project. Amongst all S protein mutations, D614G had the highest (94 %) mutation frequency. Mutations in B.1.1.7 (including 69-70del, Y144del, N501Y, A570D, T716I, S982A, D1118H and D614G) had a mutation frequency of around 5%. In contrast, mutations in B.1.351 (D80A, D215G, 242-244del, R246I, K417N, E484K, N501Y, D614G, and A701V) had a much lower mutation frequency (0.2%) (Fig.5A).

The change of binding activity between IgGs and mutant S1 was evaluated by ELISA. The change of binding activity was defined by the value of OD450_mutant S1_ / OD450 _S1_. F61, F163,C25, H184 and B110 showed low sensitivity against single-residue variants with A475V and S477I. While, antibodies H121, C25, H184 and B110 exhibited a low sensitivity towards N354K, A348T and A435S. Moreover, A199 exhibited a low sensitivity towards R246A and P384L (Fig. 5B)

**Figure 5.**
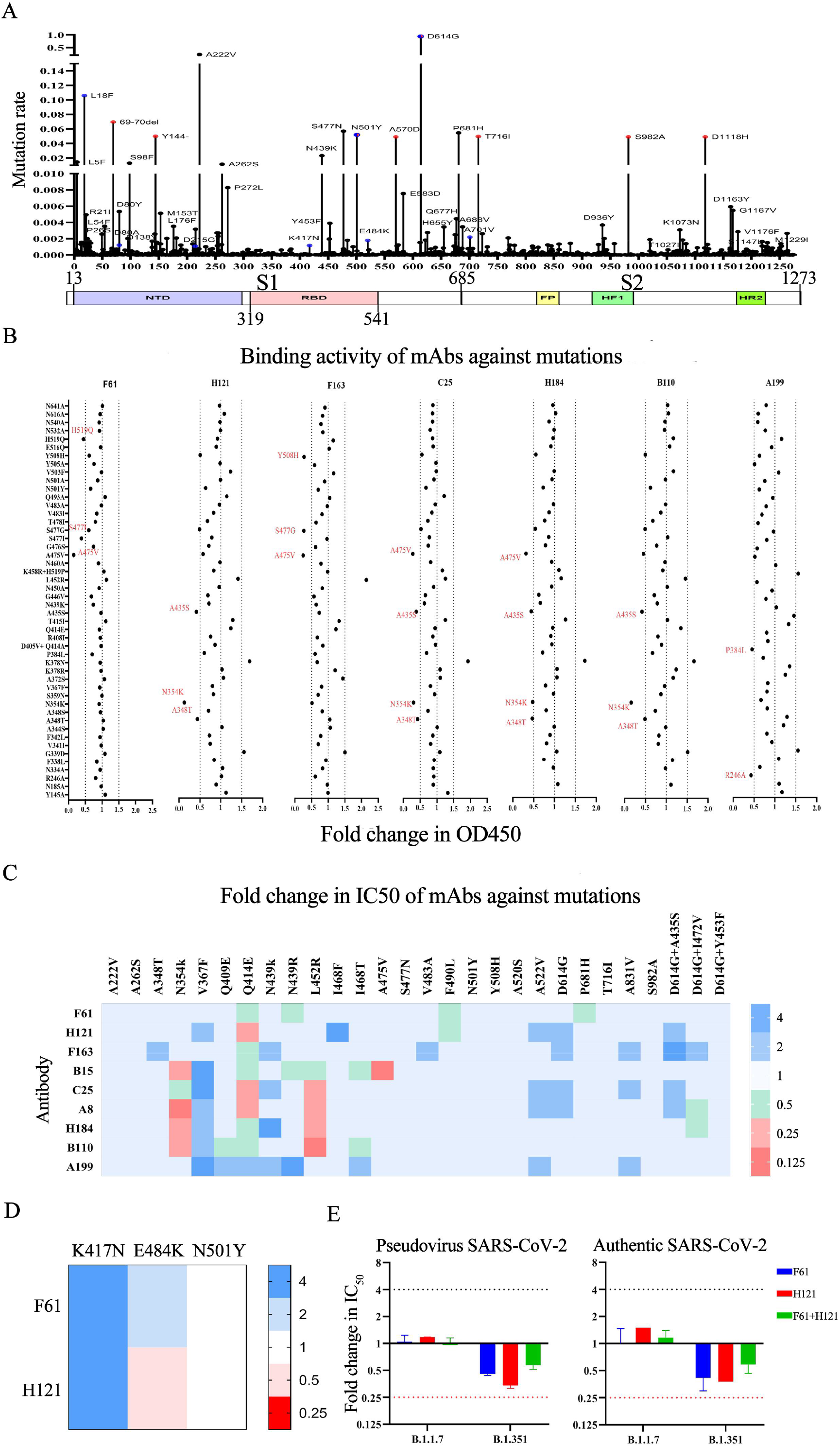
Neutralization mutations of SARS-CoV-2 S protein and their effect on antibody neutralizing activity. (A) Amino acid mutations on S protein. Mutations in B.1.1.7 lineage were labeled red. Mutations in B.1.351 lineage were labeled blue. (B)The binding activity of RBD-specific IgGs between mutant S1 protein and wild-type (WT) S1 protein was detected by ELISA. The change of binding activity was defined by the ratio of OD450mutant S1 / OD450WT S1. The dashed line indicated that the ratio was less than 0.5 or more than 1.5. The significant changes were marked red for decreased. (C)Neutralization activities of nine RBD-specific IgGs towards mutations on S protein were measured by pseudovirus. The changes in neutralization activity was showed in the ratio of IC50 between the variant and SARS-CoV-2 (GenBank: MN908947). The changes were marked with colored symbols, red for decreased, blue for increased. (D)Neutralization activities of F61 and H121 towards mutations K417N, E484K, and N501Y on S protein were measured by pseudovirus. The changes in neutralization activity was showed in the ratio of IC50 between the variant and the SARS-CoV-2 (GenBank: MN908947). The changes were marked with colored symbols, red for decreased, blue for increased.(E)Neutralization activities of F61 and C121 towards B1.1.7 and B1.351 were measured by pseudovirus and authentic SARS-CoV-2. The y-axis represents the value of IC50. The dashed line indicated that the fold change of IC50 was less than 0.25 (decrease) (marked red) or more than 4 (increase).

The neutralizing activity of mAbs against pseudovirus with mutant S protein was evaluated via neutralization assay. The change of neutralizing activity was defined by the value of IC50_SARS-CoV-2_ / IC50 _variant SARS-CoV-2_. The ratio less than 0.25 was deemed as significantly. Remarkably, F61 and F163 efficiently neutralized multiple mutations within RBD. F61 and F163 exhibited equivalent neutralization sensitivity against single-residue variant A475V and S477I, which reduced the binding sensitivity of F61 and F163. Contrarily, H121, C25, A8 were evaluated to have low potencies against Q414E. Similarly, the neutralization sensitivity of C25, A8, H184, and B110 was reduced by L452R and N354K(Fig.5C). Besides, F61 and H121 exhibited constant neutralizing activities against mutations in B.1.1.7 and B.1.351 (K417N, E484K, N501Y) (Fig.5D). Furthermore, F61 and H121 exhibited efficient neutralizing activity against B.1.1.7. F61 and H121 showed slightly decreased neutralizing activity against B.1.351, which was still efficient. Synergistic of F61 and H121 exhibited similar neutralizing activities against the B.1.1.7, B.1.351 and WT virus (Fig.5E)

## Discussion

In this study, 12 SARS-CoV-2 specific IgGs were generated utilizing the Fab phage antibody library technique which was more efficient than scFv phage library. Among the 12 selected antibodies, two of the RBD-specific antibodies (F61 and H121) had demonstrated high neutralizing activity and high affinity against SARS-CoV-2. However, one RBD-specific antibody failed to neutralize SARS-CoV-2 Meanwhile, our results from competitive ELISA assay and computer docking reviled that F61 identified a linear epitope located in residues G446-S494, which overlapped with ACE2 binding sites and H121 recognized a conformational epitope located on the sideface of RBD which not overlap with ACE2 binding sites. F61 and H121 maintained neutralizing activity against B.1.1.7 and showed a slightly decreased but still efficient neutralization against B.1.351. Synergistic of F61 and H121 exhibited higher neutralizing activity against B.1.1.7 and B.1.351. Multiple mutations on RBD could be efficiently neutralized by F61 and H121. Hence F61 and H121 had broad neutralizing activity towards SARS-CoV-2 variants.

In this study, nine high affinity RBD-specific IgGs exhibited different neutralizing activity. Two of them (F61 and H121) showed high neutralizing activity against SARS-CoV-2. However, A199 failed to neutralize SARS-CoV-2, which suggested not all RBD-specific antibodies were neutralizing antibodies. Meanwhile, our results from competitive ELISA assay reviled that F61, F163 and B15 identified ACE2 competitive epitopes. However, H121, C25, A8, H184 and B110 recognized epitopes which did not overlap with ACE2 binding sites. A199 bound to a non-neutralization epitope that did not overlap withremote from ACE2 binding sites. Results above suggested that neutralizing capacities of antibodies did not rely on their ability to block RBD-ACE2 interaction. RBD had three kinds of epitopes, thus, ACE2 competitive neutralization epitopes, ACE2 non-competitive neutralization epitopes, and non-neutralization epitopes.

Multiple mutations on S protein showed no resistant to F61 and H121. Single-residue variants(K417N, Q493A, and N501Y)on essential residues within RBM for ACE2 binding (Lan *et al*., 2020; Wan *et al*., 2020) and B.1.1.7 related mutations (N501Y, P681H, T716I and S982A) could be efficiently neutralized by F61 and H121. Most of these mutations positioned distant from target residues of F61 and H121. Therefore, the neutralization sensitivity of F61 and H121 was barely altered by them. K417N increased the neutralizing activity of F61 and H121. Replacement of the lysine with a shorter asparagine in K417N(Laffeber *et al*., 2021) increase the probability of conversion (Li *et al*., 2021) to the open conformation (Li *et al*., 2021) and destroy the salt-bridge between K417 and residue D30 on ACE2. The potential conversion in S protein trimer conformations might advance to the neutralizing activity of the two antibodies.

F61 identified a linear epitope ranging from residues G446 to S494, which inculed mutations E484K. K417N/T and E484K harbored by B.1.351 and P1 show resistance to multiple neutralizing mAbs from the first and second class (Hoffmann *et al*., 2021; Widera et *al*., 2021), including three mAbs with emergency use authorization(EUA): REGN10933 (casirivimab), LY-CoV555 (bamlanivimab), and CB6 (etesevimab)(Kuzmina *et al*., 2021; Tada *et al*., 2021; Wang L. *et al*., 2021a; Wang P. *et al*., 2021c). However, neither B.1.351 or mutations within the F61 identified epitope (A475V,L452R,V483A,Q493A) were resistant to F61. Computer docking suggested that F61 recognized a linear epitope on RBD, which was considered more stable than conformational epitope. Therefore, the single-residue mutation within F61’s epitope would barely alter the neutralizing activity of F61. Moreover, P2C-1F11(Ge *et al*., 2021) which shares a similar epitope with F61 shows no reduction in neutralizing capacity against B.1.351(Li *et al*., 2021). Compared to CB6 (Shi *et al*., 2020), another mAb that failed to neutralize B.1.351 (Li *et al*., 2021; Wang L. *et al*., 2021a) in class one, F61 and P2C-1F11 identified a linear epitope and involved a relatively high number of residues in their binding interface on RBD (Shi *et al*., 2020; Ge *et al*., 2021). Antibodies covering more critical residues involved in the RBD-ACE2 interface might have a higher tolerance to viral mutations. Moreover, viral mutations have a superposition effect on mAbs. Specifically, an increasing number of mutation sites in the RBD is correlated with the immune escape from a steadily increasing number of monoclonal antibodies (Li *et al*., 2021; Widera *et al*., 2021). Therefore, with the accumulation of mutations in RBD protein, more and more antibodies, including F61, would be escaped by SARS-CoV-2 variants.

H121 exhibited high levels of neutralization with epitopes on RBD remote from the RBM and was believed non-overlap with ACE2 binding sites. H121 might lock the S trimer in its closed state and hide the RBM region out of ACE2’s accessibility (Benton *et al*., 2020) through binding to two neighboring RBDs within an S trimer, which was found in S309 and S2M11(Pinto *et al*., 2020; Tortorici *et al*., 2020) sharing similarities with H121. Nevertheless, mutation Q414E, instead of mutations within the H121 recognized epitopes (A348T, N354K, N439K, A435S and A520S), escaped from H121. Q414E located remote from the epitope of H121 on RBD. Q414E did not reduce the binding sensitivity of H121 to RBD (Fig. 5B), determining other explanations for lower potency.

Revealed in our study, the cocktail of F61 and H121 exhibited increased neutralization to SARS-CoV-2, and showed efficient neutralization to B.1.1.7, B.1.351 and WT SARS-CoV-2. Therefore, the cocktail of F61 and H121 with broad neutralization improved treatment efficacy by mitigating viral escape (Wang N. *et al*., 2021b). F61 and H121 bond to distinct and non-overlapping regions of the RBD and masked more epitopes on RBD (Piccoli et al., 2020). Occupying more neutral epitopes prevented virus escape. Hence synergistic use of antibodies with different epitopes should be investigated for developing future therapeutic antibodies.

While designing antibody-based, efficient biological drugs, it is essential to precisely delineate the interaction between antibody and antigen structures. To achieve this, we used computer modeling and docking technique, which is more rapid and accurate compared to X-ray crystallography and cryo-electron microscopy (cryo-EM), for the binding structure prediction of antibodies and RBD. Since antibodies have a highly conserved framework, homology models building for antibodies can be reasonably accurate (Yamashita *et al*., 2014; Leem *et al*., 2016). Thus the models for F61 and H121 were highly reliable given that they shared a similarity of more than 90% with the templates we used for homology modeling. The ZDOCK method used in this study, a rigid-body docking algorithm based on fast Fourier transforms (FFTs) (Pierce *et al*., 2014), does not consider possible conformational changes, causing possible deviation in docking results. Therefore, we will consider semi-flexible docking protocols accounting for protein flexibility, such as HADDOCK (van Zundert et al., 2016), and dynamic simulation methods for further model optimizations.

Succinctly, two antibodies (F61 and H121) obtained within our study were demonstrated with high affinity and high-neutralizing activity against distinct RBD epitopes. Meanwhile, evidence generated during our screening indicated that not all RBD-specific antibodies were capable of performing neutralization. In comparison, antibodies could neutralize SARS-CoV-2 without necessarily blocking the binding of RBD to ACE-2. Neutralizing activity of F61 and H121 was mostly maintained when tested against multiple mutations and variant B.1.1.7 and B.1.351, which revealed a broad neutralizing activity against SARS-CoV-2 variants. While our findings could be further solidified with dynamic simulation methods, they provided a guideline for the current and future vaccine design, therapeutic antibody development, and antigen diagnosis of SARS-CoV-2 and its novel variants.

## Acknowledgements

We thank XL. Shi from Tsinghua University for her technical help in human Antibodies expressing; TY. Chen and his research from INNOVITA for supporting antibody detection reagents of SARS-CoV-2; HW.Wei from Jiangsu East-Mab Biomedical Technology for his advice in high-yield antibody expressing; W.An for his technical help in performing SPR. This work was supported by the National Science and Technology Major Project (2018ZX10711-001)(2016ZX10004222-002).

## Author Contributions

YY. Qu performed the experiments and wrote the paper; LA. Sun, MY. Wang, and performed the experiments; YZ. Jing, FX. Zhan and YC. Wang contributed reagents/materials/analysis tools. W. Wei, QL. Yin, MF. Liang, Y. Lang, JD. Li, C. Li and DX. Li analyzed and discussed the data. MF.Liang, SW.Wang and WX.Guan designed the project and edited the manuscript. All authors read and approved the final manuscript.

## COMPLIANCE WITH ETHICS GUIDELINES

### Animal and Human Rights Statement

This article does not contain any studies with human or animal subjects performed by any of the authors.

---

**Figure S1.**
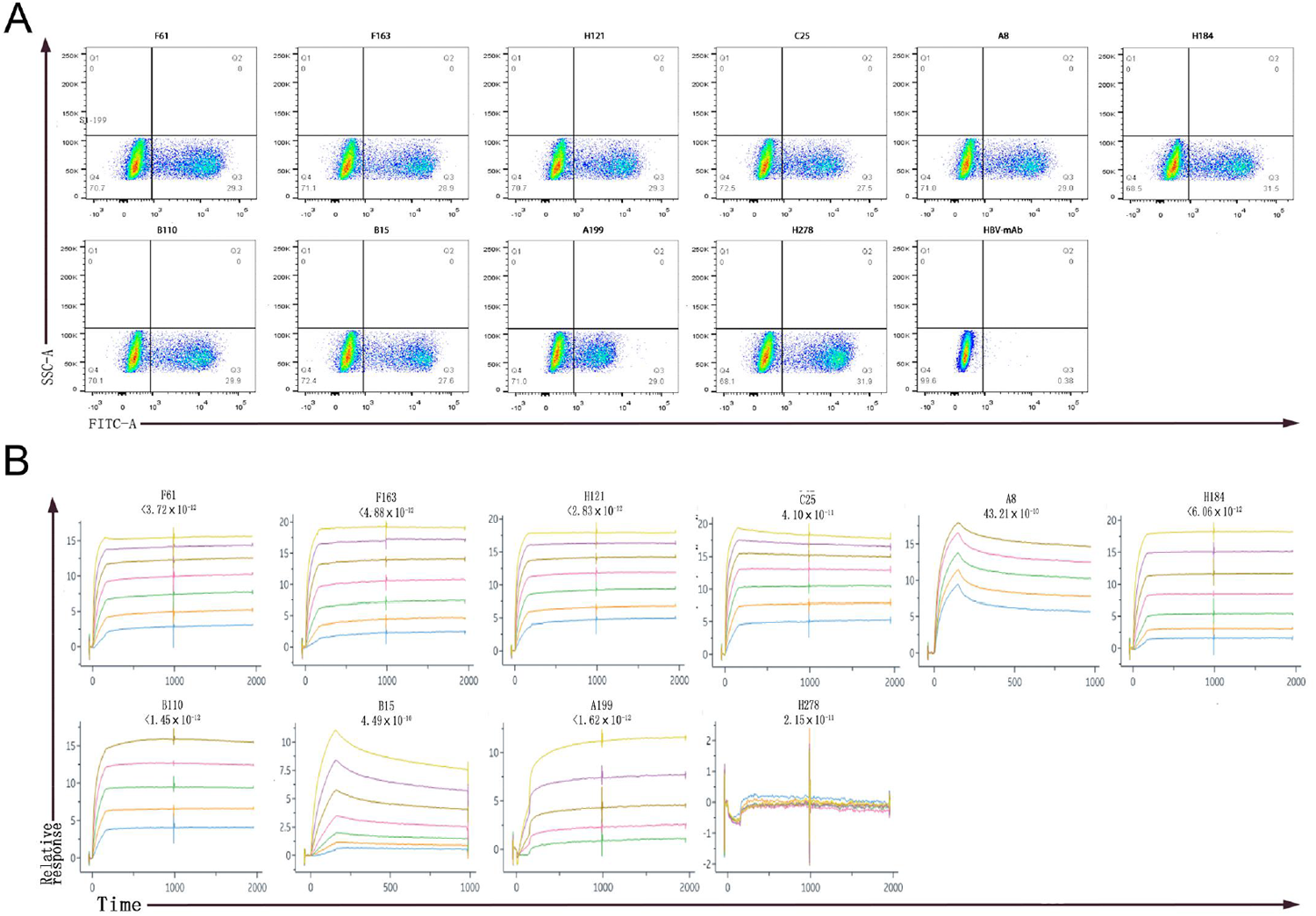
Characterization of candidates IgGs.(A) The specificity of candidates antibodies detected by FACS. HEK 293T cells expressing SARS-CoV-2 S protein were incubated with candidate antibodies or isotype IgG (HBV mAb) and then stained with anti-human IgG FITC-conjugated antibody. The X-axis represented the fluorescence intensity of human antibodies labeled by FITC. (B) The affinity between antibodies and S1 was measured by BIAcore 8000 system.

